# Phylogenetic Studies and Inhibitor Design Targeting Protein Interacting Interface of Nucleoid-Associated Protein HU

**DOI:** 10.1101/2020.06.18.135426

**Authors:** Debayan Dey, Suryanarayanarao Ramakumar

**Author notes:** To whom correspondence should be addressed: Debayan Dey: Department of Biochemistry, Emory University School of Medicine, 1510 Clifton Road NE, Atlanta, GA, 30322.

## Abstract

The formations of nucleoprotein structures by promiscuous DNA binding proteins like HU are assisted with their protein protein interaction capability with other proteins. In *E. coli* Gal repressosome assembly, GalR piggybacks HU to the critical position on the DNA (hbs site) through a specific GalR–HU interaction using an interface at the bottom of alpha helical region, which we termed as HUpb interface. Similarly, MtbHU also interact with Topoisomerase I with the same interface to enhance its relaxation activity. In an earlier study, we determined the crystal structure of MtbHU, inhibited it using stilbene derivatives which inhibited the cell growth. It motivated us to understand the evolutionary and structural characteristics of the HUpb interface, which has not been investigated previously for HU or for any other NAPs. Our analyses found residue positions corresponding to MtbHU Thr11 to Gln20 form the interface while Ala23 serves the pocket lining residue. Due to ancestral mutations in the duplication event before the HU and IHF split, physicochemical properties of the interface vary among clades. Thus, this interface could engage different proteins in different HU oligomeric states in *Proteobacteria*. Protein-protein interfaces are usually a challenging target due to its flatter surface. In case of MtbHUpb interface, we observed that due to the presence of a partially hydrophobic pocket, small molecule scaffolds could fit into it, while the ligand can be further designed to interact with D17, which is the crucial residue for Topoisomerase I interaction. We used a two-step virtual screening protocol with known drug like molecules as starting set to an aim to re-purpose drugs. Our docking results showed compounds like Maltotetraose, Valrubicin, Iodixanol, Enalkiren, indinavir, Carfilzomib, oxytetracycline, quinalizarin, Raltitrexed, Epigallocatechin and their analogues exhibit high scoring binding at MtbHUpb interface. Our present report gives a model example of an evolutionary study of an interface of nucleoid associated protein, which is used to computationally design inhibitors. This strategy could be in general useful for designing inhibitors for various types of protein-protein interfaces using evolutionary guided design.

## Introduction

Nucleoid-associated proteins (NAPs) play an important architectural role in DNA compaction as well as a regulatory role in various DNA transaction processes like replication and recombination (Dorman, 2009; Dillon and Dorman, 2010). They have the ability to alter the global conformation of DNA by bending, wrapping and bridging it. The higher order superstructures they form, influence the transcriptional landscape of the bacterial cell. The earlier notion of a single function of a single protein is presently being challenged and an increasing number of proteins are being identified as being multifunctional (Jeffery 2003). NAPs are generally believed to interact with DNA performing various architectural and regulatory functions, but they also interact with proteins through a different interface, recruiting other proteins into a nucleoprotein assembly.

Previous studies have shown the formation of nucleoprotein structures by promiscuous DNA-binding proteins which are targeted to specific DNA locations by their interaction with other DNA-binding transcription factors (Thomas and Travers 2001). HMG-1 belongs to a family of conserved chromatin-associated nucleoproteins that bend DNA and also facilitate the binding of various other DNA binding proteins to their cognate DNA sequences. HMG-1 binds specifically to p53 Tumor suppressor protein and stimulates its DNA binding (Jayaraman et al. 1998), interacts via establishing protein-protein contacts with HOX homeodomain proteins (Zappavigna et al. 1996) and Oct family transcription factors which markedly increase their sequence-specific DNA binding activity (Zwilling et al. 1995).

HU is a small dimeric NAP ubiquitously found in bacteria and plays a pivotal role in shaping the nucleoid structure (Dey 2020). HU and Integration Host Factor (IHF) belong to prokaryotic DNA-bending protein family II (DNABII protein) and consist of three alpha helices and five beta strands, where the beta strands from each protomer form the DNA binding cradle while the alpha helices form the dimerization interface representing the HU/IHF fold. These proteins are involved in cell division (Dri et al., 1991) replication (Ryan et al., 2002), DNA recombination and repair (Kamashev and Rouviere-Yaniv, 2000), DNA supercoiling (Bensaid et al., 1996) and CRISPR-Cas mediated integration (Nuñez et al., 2016).

In E. coli Gal repressosome assembly, a DNA loop is formed by HU to facilitate the GalR–GalR interaction, separated by to two distal operators. It was reported that during this nucleoprotein complex formation, GalR piggybacks HU to the critical position on the DNA (hbs site) through a specific GalR–HU interaction (Kar and Adhya 2001). The site of HU which interacts with GalR is composed of an interface formed between two alpha helices (α1 and α2) near the N-terminal region. The interface is contributed from both the protomer chains. Similar results of a direct regulation of Topoisomerase I by M. tuberculosis HU (MtbHU) was recently reported, to enhance its relaxation activity, in which MtbHU interacts with Topoisomerase I utilizing the same interface (Ghosh et al., 2015). Interestingly other than primary function of DNA binding, HU is also associated with other moonlighting functions which include adhesion like function at the extracellular surface of Mtb, while interacting with antigen85 complex (Katsube et al. 2007). Moreover, in many organisms HU like proteins are essential, which makes them a promising drug target for infectious pathogens, which encouraged our group to design structure-based inhibitors against them. In an earlier study, our group has determined the crystal structure of MtbHU, inhibited it using stilbene derivatives which curtailed the cell growth (Bhowmick et al.,2014).

The previous studies motivated us to understand the evolutionary and structural characteristics of the HU protein binding interface (HUpb interface) (Fig.1a,b), which has not been investigated previously for HU or for any other NAPs. In our previous work, we discussed the phylogenetic divisions in HU/IHF proteins and the sequence and structure based determinants of their (multi-)specificity (Dey et al. 2017). In the present investigation, we have analyzed the evolutionary and structural makeup of HUpb interface across species. The structural features include formation of cavities or pockets with different physicochemical makeup, which influences their partner binding preferences. We performed a maximum likelihood based phylogenetic analysis on 117 HU and IHF sequences from well sampled pathogenic and model prokaryotes. We also performed ancestral protein reconstruction aimed at understanding the evolution of HUpb interface. As a case study, we performed a systematic virtual screening, molecular docking and chemoinformatics study for finding MtbHUpb inhibitor.

**Fig. 1:**
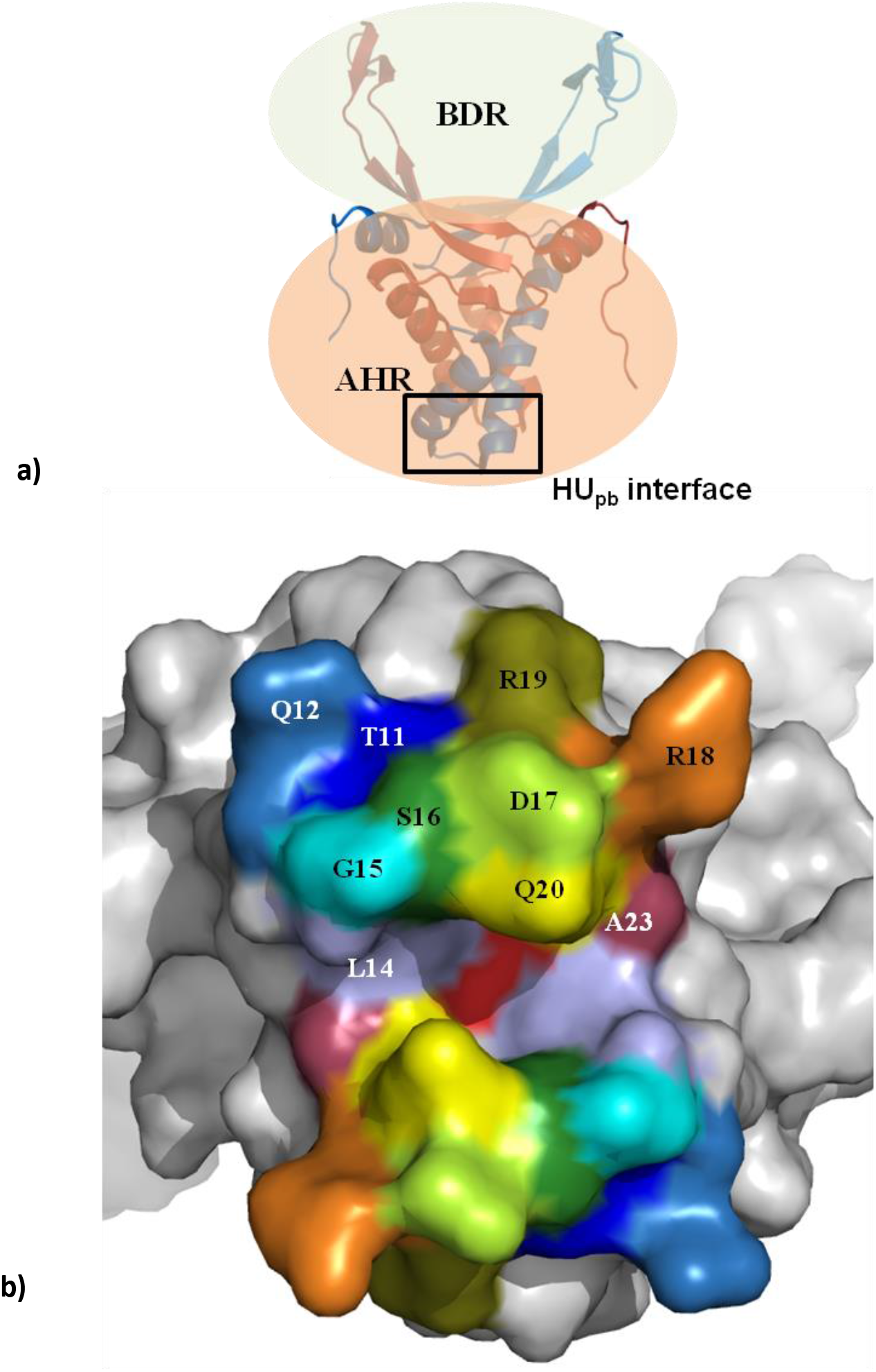
**a)** HU structure can be segregated into two segments, alpha helical region (AHR) which forms the dimerization core, interface for DNA stabilization (DNA draping region) and HU_pb_ interface which is involved in interaction with other proteins and beta-sheet region (BDR) which engages in binding to DNA and bending. **b)** MtbHU_pb_ interface (from crystal structure PDB: 4PT4) is analyzed to determine the residue positions which contribute to the formation of this interface. We defined the HU_pb_ interface as the region in the vicinity of the loop connecting α1 helix to α2 helix, which is completely or partially solvent exposed. T11, Q12, L14, G15, S16, D17, R18, R19, Q20 and A23 (shown only the half part of the interface due to Non crystallography symmetry (NCS) related other half. MtbHU_pb_ Pocket is formed by L24 and A23, while D17 acts as a beacon residue for its interaction with other protein.

## Materials and Methods

### Sequence dataset construction, phylogenetic analyses and ancestral sequence reconstruction

HU/IHF proteins (InterPro ID IPR00019) were selected from diverse phyla e.g. Actinobacteria, Bacteroidetes, Firmicutes, Proteobacteria, Spirochetes, Fusobacteria and Chlamydiae. Multiple sequence alignment guided by structural information was performed by MEGA 7.0 using CLUSTALW algorithm. ProtTest 1.4 (Abascal et al., 2005) was used to determine the best-fit amino acid substitution model and parameter from values for each data set. In this case, the (Le and Gascuel 2008) LG model was the best fit according to the Akaike information criterion (AIC). PHYML 3.0 was used to run the maximum likelihood (ML) phylogenetic construction (Dey et al. 2017). The subtree pruning and regrafting method were used to search tree topology. Branch support for the resulting topology was determined by the Shimodaria–Hasegawa-like approximate likelihood ratio test. Ancestral sequence reconstruction was performed using ML method using MEGA 7.0. We further confirmed the phylogenetic analysis using evolutionary trace method to understand the specificity determining residues in a larger protein family or superfamily. This method was previously utilized in several studies to determine functional residue differences among different phylogenetic clades and compares closely to ML method with less computational calculation time(Dey 2018, Shrilakshmi et al. 2019, Nosrati et al. 2019, Kuiper et al. 2019, Dey et al. 2020).

### Structural analysis, homology modeling and pocket feature calculation

We calculated the hydrophobic, electrostatic and H-bonding interaction of the HUpb interface using PIC server (Tina et al., 2007). Homology modeling of HU and IHF homologs were performed using Swiss Modeler server, with further energy minimizations using Chimera using 1000 steps of steepest descent and 1000 steps of conjugate gradient. For pocket analysis, PISA and DoG Site Scorer server were used. Figures were generated using PyMOL (deLano Scientific).

### Virtual screening, chemoinformatics and docking

The protein structure of MtbHU was taken from the Protein Data Bank (PDB) with PDB ID code 4PT4. The Protein Preparation module on Maestro with default settings was used to prepare the protein structures. 3D conformations were generated for each compound by LigPrep, (Schrodinger, LLC, New York, NY, 2013) with default settings with OPLS2005 force field. A two-step virtual screen was performed against MtbHUpb pocket using DrugBank dataset first to identify existing drug like chemical scaffolds, which was further used a guide set for a larger virtual screening on Chembridge and PubChem libraries. We used the chemoinformatics module Canvas for calculation of ligand properties and hierarchical clustering. The clustering were performed based on Daylight’s Fingerprint and Tanimoto similarity. Docking (SP and XP) was carried out using Glide (Schrodinger,LLC, New York, NY, 2013) on Maestro version 9.6. A docking grid of 20 Å3 was created around the MtbHUpb pocket with centroid of Ala 23 from both chains as centre of docking grid. In later docking steps we focused on ligands with high score as well as shielding Asp17.

## Results and discussions

### Phylogenetic study and structural variability of HUpb interface residues

In our previous work (Dey et al. 2017), we analyzed the phylogenetic distribution of HU/IHF proteins and determined the factors responsible for their DNA binding sequence specificity (Dey et al. 2017). In the present study, we aligned the 117 amino acid sequences (distributed across various phyla), and then subjected them to phylogenetic analysis using maximum likelihood (see Methods). The phylogenetic tree was majorly partitioned into HU, IHFα and IHFβ, similar to results in the Dey et al. 2017 (Fig. 2). In the present dataset, we included HU like proteins from eukaryotes and viruses to understand their origin from various bacterial phyla and also to examine their physicochemical and structural features of HUpb interface residues. HU like protein encoded in the plasmid of Red Algae Galdieria sulphuraria was found to be phylogenetically related to Cyanobacteria. Interestingly, HU from swine flu virus was found to be related to its homolog in Coxiella genus. Intriguingly, HU like proteins from Plasmodium falciparum (PfHU), Toxoplasma gondi and Staphylococcus phage, is more closer to IHFα clade.

**Fig. 2:**
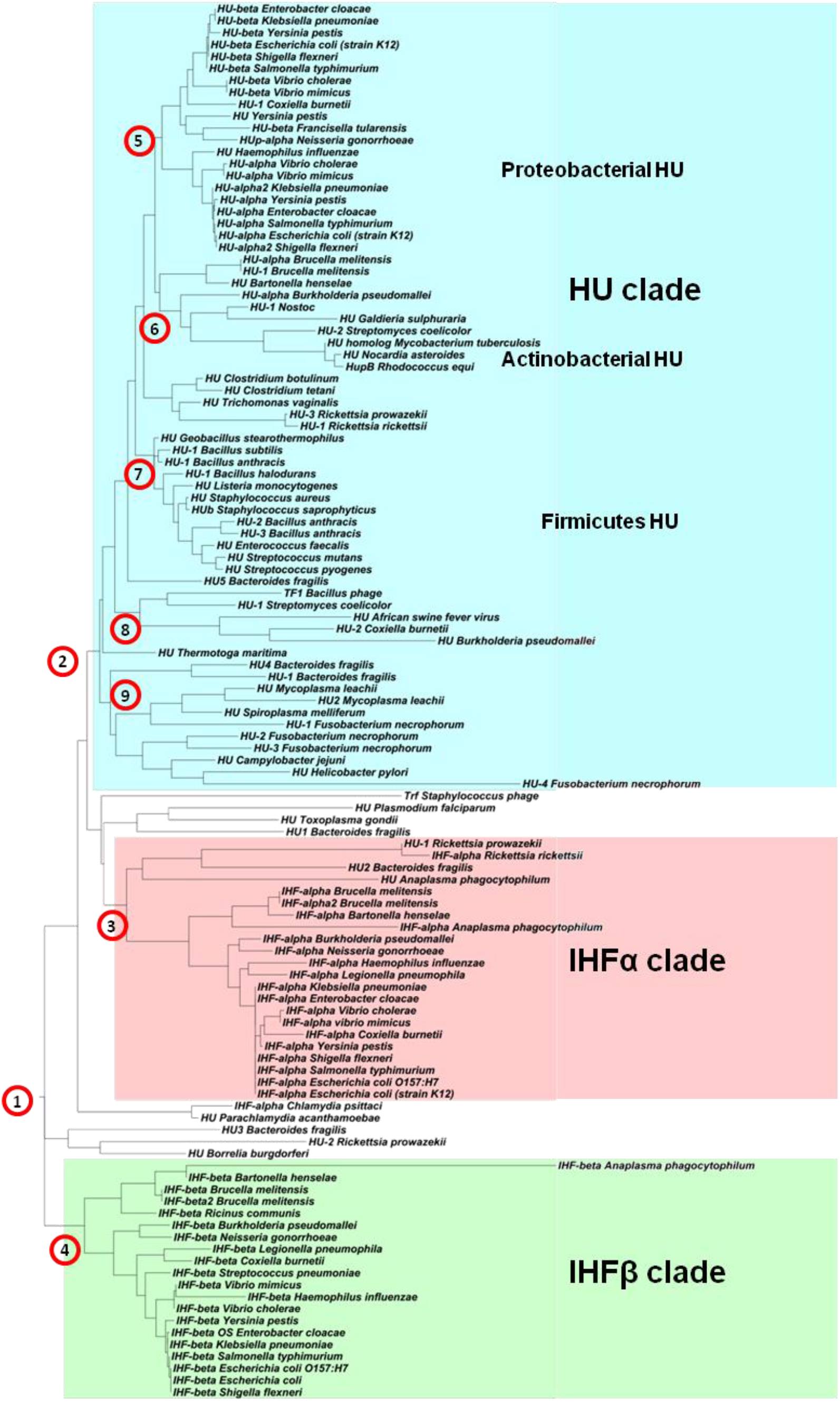
Ancestral states were inferred using the Maximum Likelihood method under the Le_Gascuel_2008 model. The tree shows a set of possible amino acids (states) at each ancestral node based on their inferred likelihood. The initial tree was inferred using the method. The rates among sites were treated as a Gamma distribution with invariant sites using 0 Gamma Categories (Gamma with invariant sites option). The analysis involved 117 amino acid sequences. All positions with less than 95% site coverage were eliminated. That is, fewer than 5% alignment gaps, missing data, and ambiguous bases were allowed at any position. The phylogenetic tree is majorly partitioned into three clades HU (blue), IHFα (red) and IHFβ (green). We also observe in the HU clade, smaller phyla specifc sub-clades are also present which shows differences in the HU_pb_ interface. Evolutionary analyses were conducted in MEGA7. The phylogenetic tree is majorly partitioned into three clades HU (Blue), IHFα (Red) and IHFβ (green), with various other sub-clades in each. HU clade is observed to be segmented into *Proteobacterial*, *Actinobacteria*l and *Firmicutes* sub-clades. The numbers in red circles represent ancestral nodes of various clade and sub-clade, with root ancestral node 1.

HU - IHF proteins structure can be segregated into two segments, alpha helical region (AHR) which forms the dimerization core, interface for DNA stabilization (DNA draping region) and HUpb interface which is involved in interaction with other proteins and beta-sheet region (BDR) which engages in binding to DNA and bending it (Fig 1a). Our inspection of HUpb interface residues of three dimensional structures from HU-IHF family proteins revealed the presence of pocket in two structures, MtbHU (PDB 4PT4) and Thermotoga maritima HU (PDB 1B8Z). MtbHUpb interface has a pocket of volume 229.7Å3, surface area 503.2Å2 and depth of 10.3Å (Fig 1b, Table 1). We defined the HUpb interface as the region in the vicinity of the loop connecting α1 helix to α2 helix, which is solvent exposed. Our analyses found residue positions corresponding to MtbHU Thr11 to Gln20 forms the interface while Ala23 serves the pocket lining residue (Fig 1b). Thus, we focused on these sites to explore the clade specific variability and structural features, which would ultimately determine its binding partner’s affinity and specificity. We also observed Thermotoga maritima HU also has a pocket of volume 170.6Å3, surface area 378.3Å2 and depth of 10.18Å, which is a smaller pocket in comparison to MtbHU’s pocket (Table 1).

**Table 1:**
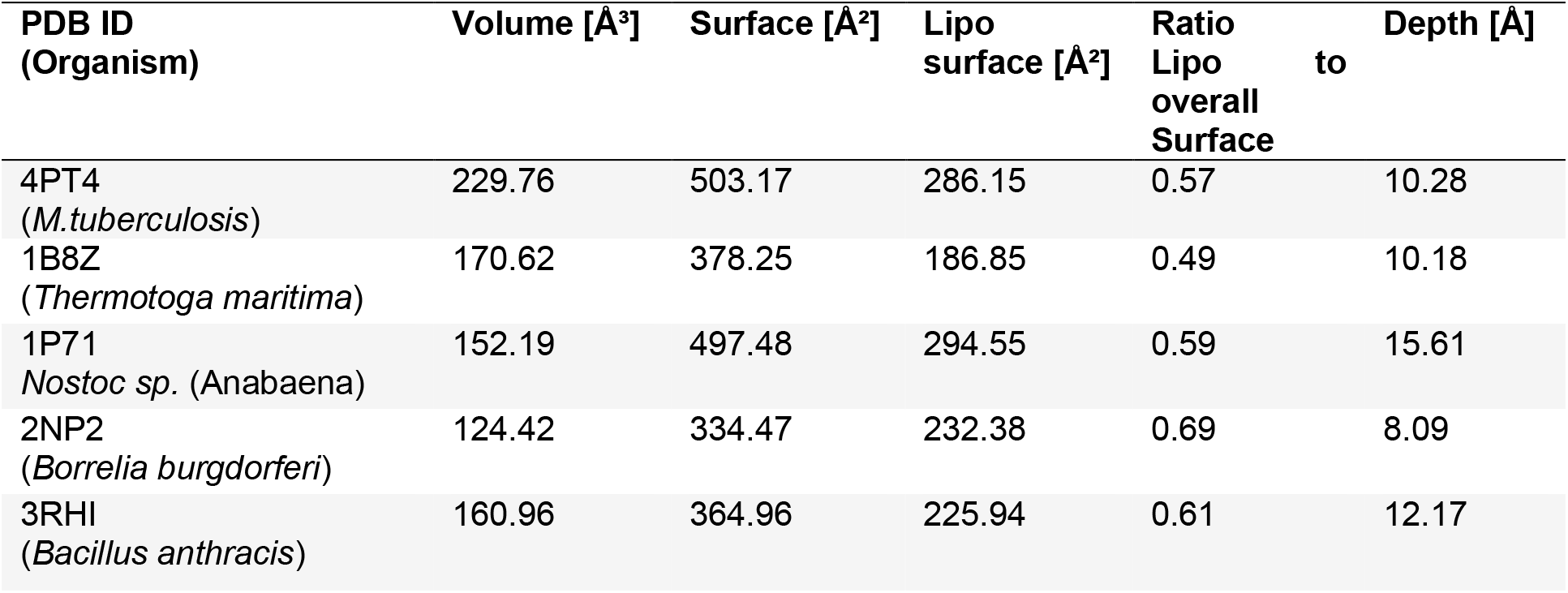
The pocket features of HU bottom region in various homologs.

Comparison of MtbHUpb interface with that in other structures showed a flatter interface, but with different electrostatic or physicochemical properties (Table 1). Although, both type of interface, one with pocket (MtbHU) and other without pocket (E. coli HU) can engage in protein mediated interaction (Kar and Adhya 2001, Ghosh et al. 2015), we speculate that the formation of pocket is species or clade specific feature and not a prerequisite for its protein binding ability. The variations among the HUpb interface residues indicate their “evolutionary mouldability” towards co-evolution with other proteins. But, there an advantage in targeting HUpb interface with pocket rather than a flatter interface, as the pocket can serve as an anchor in holding core ligand moiety, while other part of the ligand can shield crucial protein interacting hot-spot residues on the surface. Thus, in the present study, we targeted the MtbHUpb interface to computationally discover inhibitory compounds which can disrupt its interaction with Topoisomerase I. Further studies of experimental nature would be required to analyze the effect of these inhibitory compounds in the cell.

While analyzing the residue level differences in E. coli HUpb (considering HUαα homodimer for comparison purposes) and MtbHUpb interface, we observed that a single substitution of polar residue at 16th position determines the presence or absence of pocket. In MtbHU, this position is occupied by Ser, while in E. coli (and other Proteobacterial and Firmicutes HU) this position is occupied by Leu (or other hydrophobic residues). Our structural alignment of E. coli HUpb and MtbHUpb interface residues resulted in a low R.M.S.D among the main chains (less than 0.5 Å), where Leu16 from both the protomers of E. coli HU draws closer (Cβ distance 7.9 Å) due to hydrophobic effect (Fig 3a, c) and Ser16 from both the protomers of MtbHU are separated away (Cβ distance 9.2 Å), interacting with solvent, thus creating a pocket in the HUpb interface (Fig 3a, b). Confirming our result, we found similar pocket is formed in Thermotoga maritima HU also, where the position is occupied by Ala, which due to smaller size, could not fill up the gap, thus creating the pocket (Fig 3d, e). We also observed the interface formed by IHFαβ heterodimer, which is asymmetric in nature and shows a small depression near IHFβ chain’s Ile17. Similar to E. coli HU, position 16th in IHFα is occupied by Leu18 and in IHFβ Ile17, which draws closer due to hydrophobic effect. But, due to the branched nature of hydrophobic side chain in Ile, it is shorter than Leu (Leu 18 of IHFα) to fill the gap, thus creating a depression (Fig.4 a-c). This further reiterates the role of position 16 in formation of pocket in the HUpb interface.

**Fig. 3:**
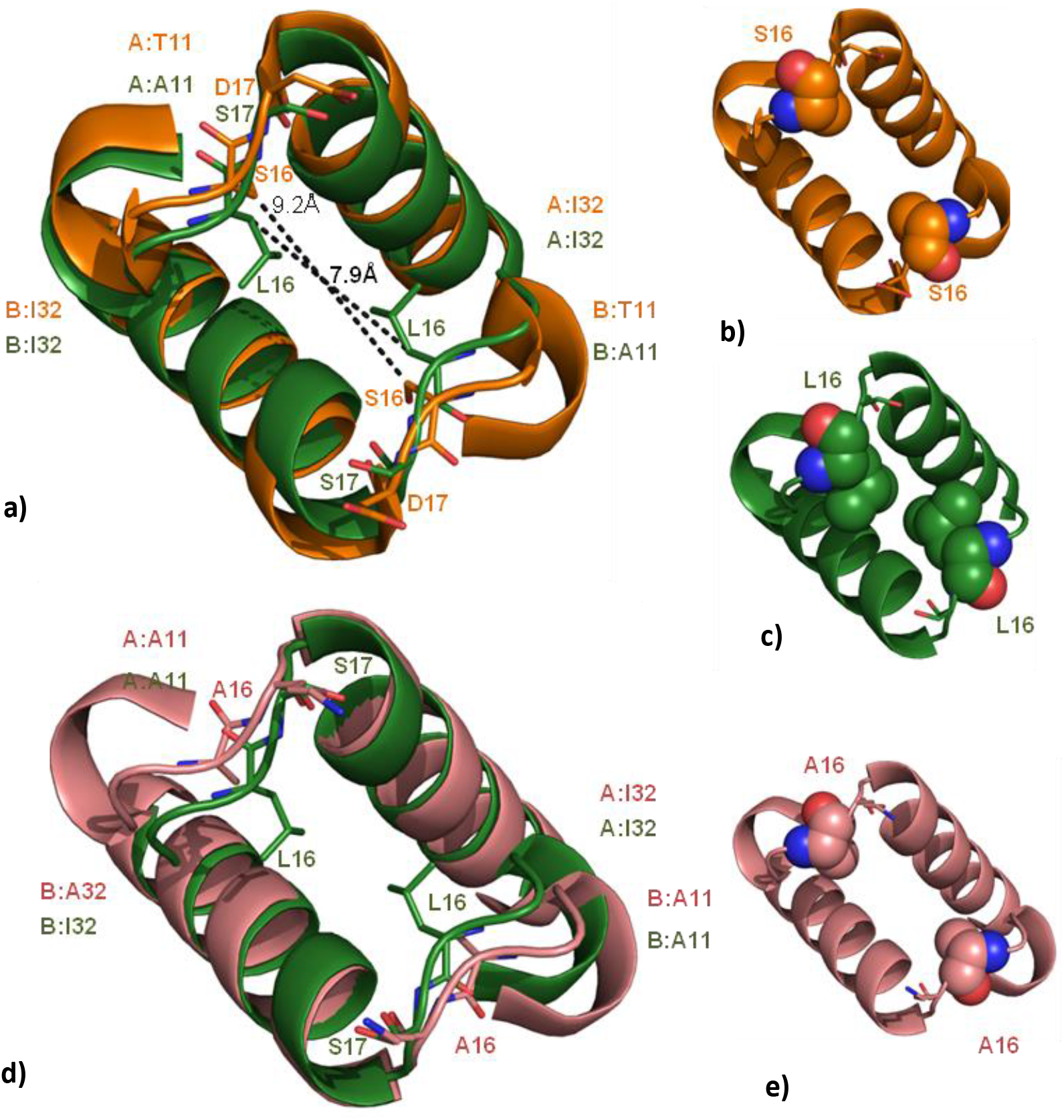
Effect of a single residue position in formation of pocket at HU_pb_ interface. **a)** Structural superimposition of region forming HU_pb_ interface between *E. coli* HU_αα_ (PDB 1MUL, shown in green) and MtbHU (PDB 4PT4, shown in orange). Although R.M.S.D is very low among the main chains (less than 0.5 Å), key side chains residues play crucial role in shaping its interface ruggedness. Position 16 in *E. coli* HU_αα_ is occupied by Leu which draws inwards with hydrophobic interactions (C_β_ distance 7.9 Å) while in MtbHU, Ser residue at this positions drive outwards (C_β_ distance 9.2 Å) interacting with solvent, thus creating a pocket in the HU_pb_ interface. A comparison of van der Waals surface of S16 from each protomer of MtbHU **(b)** and L16 of each protomer of *E. coli* HU_αα_ **(c)** is shown. **d))** Structural superimposition of region forming HU_pb_ interface between *E. coli* HU_αα_ (PDB 1MUL, shown in green) and *Thermotoga Maritima* HU (PDB 1B8Z, light pink), where the same position is occupied by Ala, which also creates a pocket in the HU_pb_ interface. e) van der Waals surface of A16 from each protomer *Thermotoga Maritima* HU is shown.

**Fig. 4:**
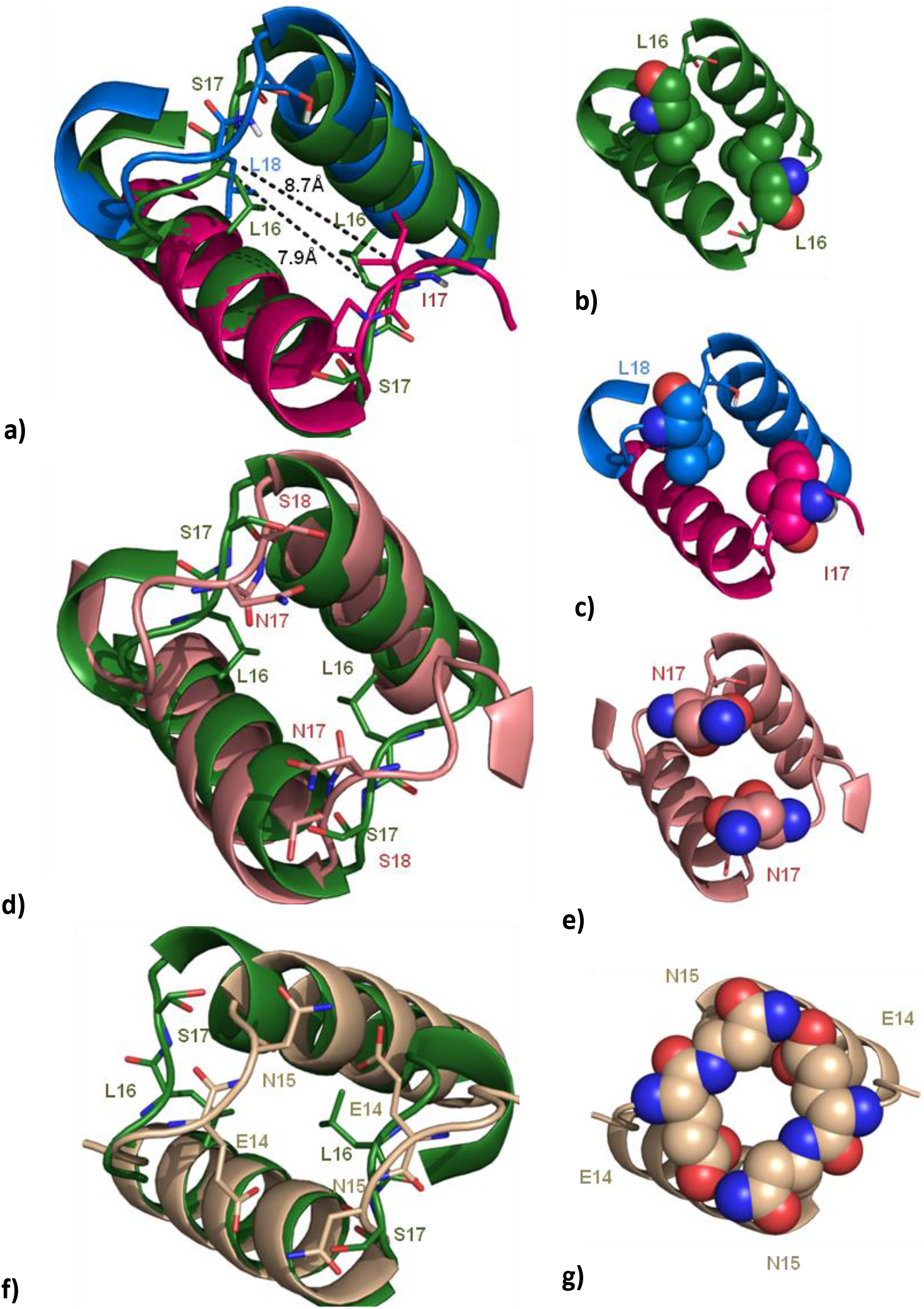
Effect of a single residue position in formation of pocket at HU_pb_ interface. **a)** Structural superimposition of region forming HU_pb_ interface between *E. coli* HU_αα_ (PDB 1MUL, shown in green) and *E. coli* IHF (PDB 1IHF, chain A and B shown in blue and pink respectively). In the 16^th^ position of our alignment, we observe that IHFα has L18, while IHFβ has I17, which draws close due to hydrophobic interaction (C_β_ distance 8.7 Å). Compared to *E. coli* HU_αα_ L18 interactions shown in **b)** smaller side chain of Ile, it still leaves a gap, shown as **c)** van der Waals surface of L18 and I17 from each chains. **d)** Structural superimposition of region forming HU_pb_ interface between *E. coli* HU_αα_(PDB 1MUL, shown in green) and *Helicobacter Pylori* HU shown in light pink color. Is a modeled structure which shows presence of polar residue like Asn in 16^th^ position, leading to similar pocket forming feature as MtbHU. **e)** van der Waals surface of N17 from each protomer of *Helicobacter Pylori* HU is shown. **f)** Structural superimposition of region forming HU_pb_ interface between *E. coli* HU_αα_(PDB 1MUL, shown in green) and *Plasmodium falciparum* HU shown in light brown color is a modeled structure which shows presence of negatively charged residue like Glu (E14) in 16^th^ position, leading to similar pocket forming feature as MtbHU**. g)** van der Waals surface of E14 and N15 from each protomer of *Helicobacter Pylori* HU is shown, which shows it forms a pore like structure in the HU_pb_ interface.

To further understand the role of this position, we homology modeled two HU homologs from Helicobacter pylori and Plasmodium falciparum. The choices of these two proteins were made from phylogenetic inferences (Fig. 5.1). We observed that although mostly residue position 16 is occupied by hydrophobic residues, but in Actinobacterial HU and a small branch consisting HU homologs from Helicobacter pylori, Fusobacterium necrophilum and Campylobacter jejuni, this position is occupied by polar residues Ser and Thr. Also, the HU homolog from Plasmodium falciparum shows Glu at this position. Our structural alignment of E. coli HU and both Helicobacter pylori and Plasmodium falciparum showed the formation of pocket in the latter (Fig.4 d-g). Polar or negatively charged residues at this position are not drawn together as in case of hydrophobic residues, thus creating a pocket. In case of Plasmodium falciparum HU, we observed the modeled structure could possibly form a pore formed by Glu14 and Asn15 from both the protomers (Fig.4g). Taken together, these data reveal that polarity of position 16 is determinant in formation of pocket in HUpb interface.

**Fig. 5:**
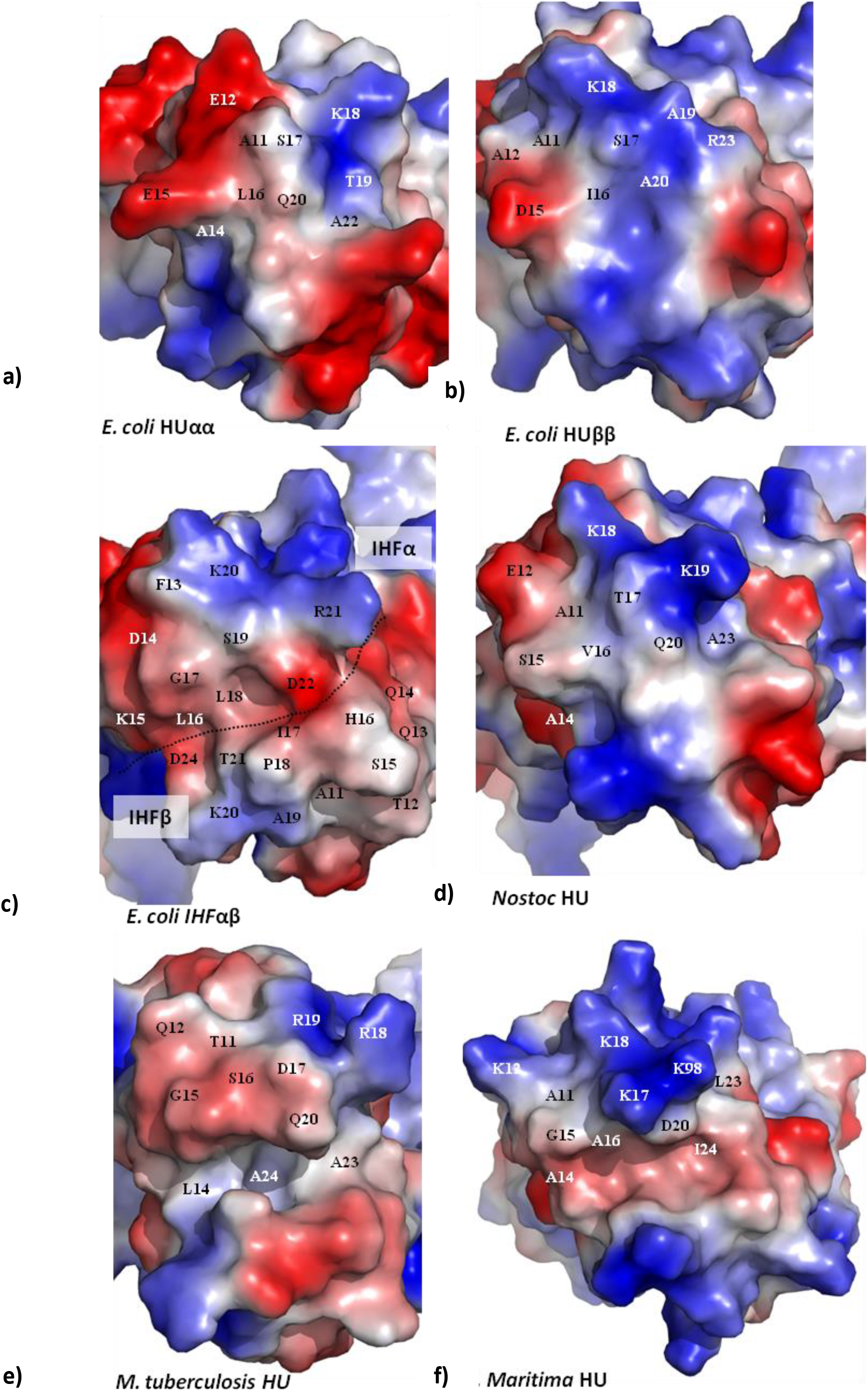
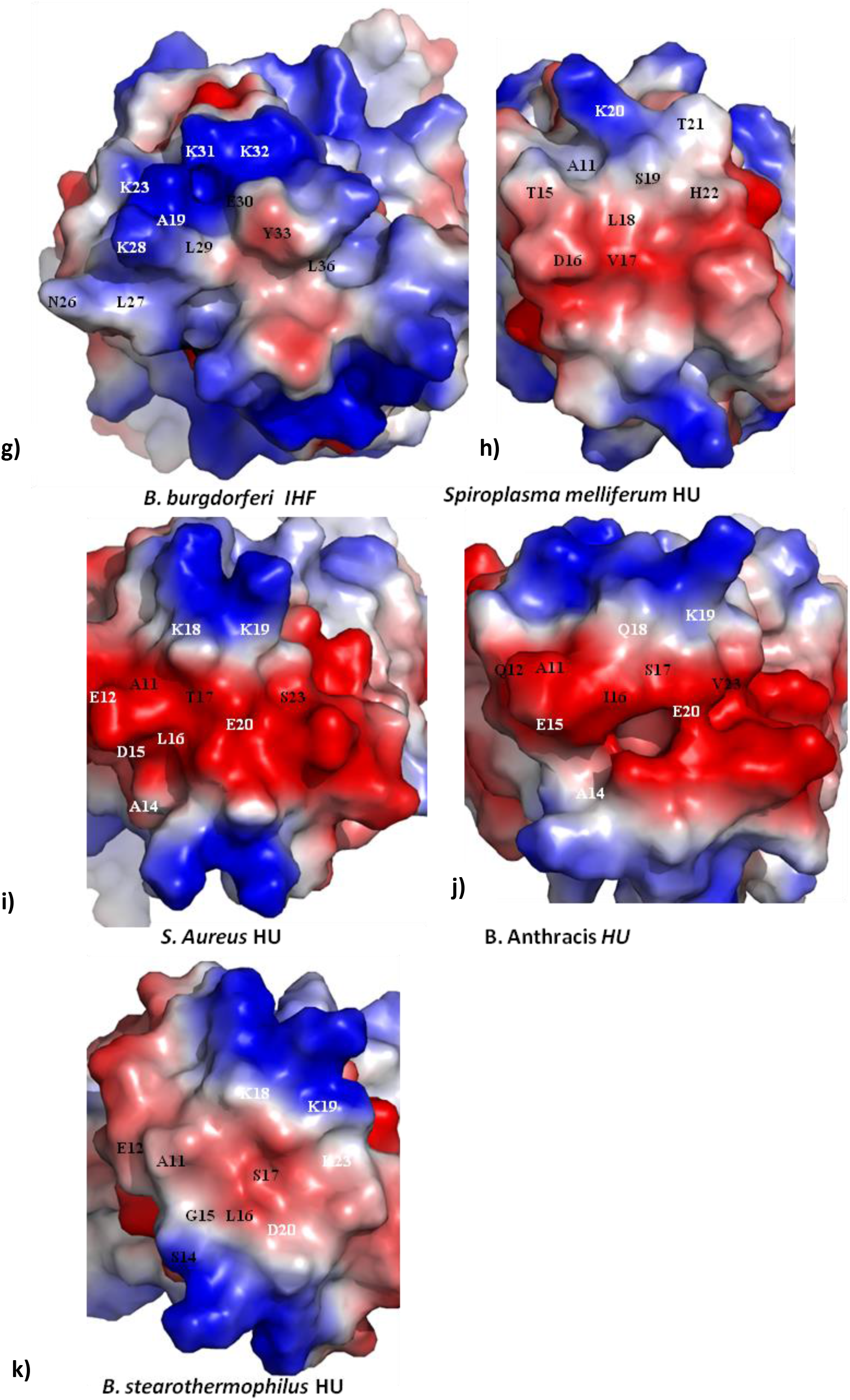
**a) –k)** are the surface electrostatic potential of HU_pb_ interface for various crystal structures showing residues on the surface as well as in the pocket. Except IHF, all the proteins shown here are homodimeric so labeling is done only for one of the protomer as other half part of the interface is related by Non crystallography symmetry (NCS). We observe a wide diversity of surface ruggedness as well charge distribution, which influence the nature of its protein partner interacting via this interface.

Protein-protein interaction depends on the shape as well as the physicochemical properties of the interface. Thus, the evolution of residues on the interface directly plays role in determining its binding partner. In most of the cases, the evolution of interface residues of the two binding partners are correlated (Goh et al. 2000). We observed differences interface of homo and heterodimer of E. coli. HUαα is more negatively charged compared to HUββ interface (Table 1, Fig 5 a,b). Differences also lies in the position 19 and 20, wherein HUββ it is Ala, while in HUαα it is polar in nature (Table 1). The conformation flexibility in formation of HUpb interface is observed while comparing the solvent assessable surface area of HUpb interface residues in the same protein in different oligomerization states. We observed that HUα chain in homodimeric form (PDB: 1MUL) and in heterodimeric form (PDB: 2O97) shows large differences in residue solvent assessability in Glu12, Ala14 and Lys18 (Table 1). Similarly analyzing the E. coli IHF heterodimer showed predominance in negatively charged residues in one half (Asp14, Asp22 and Glu25 in IHFα chain) and enrichment of polar residues in the other half (Thr12, Gln14, Ser15, Pro18 and Thr21) (Table 1 and Fig. 5c). Thus, our data points that, even in the same organism, the differences in the dimerization type changes the interface which could engage in interactions with different proteins.

In Borrelia burgdorferi Hbha, a predominant positive charged patch contributed by Lys in 12, 18 and 19 positions (Fig 5g). We also observed that HU homologs from Firmicutes e.g. S. aureus, B. anthraces and B. stearothermophilus are composed of a majority of negatively charged residues at the HUpb interface contributed by residues in position 12, 15 and 20 (Table 1, Fig. 5i-k). Although various physicochemical differences in HUpb interfaces residue positions, we observed that the beacon or hot-spot residue position, which interacts with GalR in E. coli (Ser17) and Topoisomerase I in Mtb (Asp17) (Kar and Adhya 2001, Ghosh et al. 2015) is polar or negatively charged in nature.

Also, we observed that residue position 18 and 19, which also contributes to protein mediated interactions (Kar and Adhya 2001), is conserved with positively charged residues. We also observed crystal structures in which E. coli HU dimer (PDB ID: 4YF0) exhibits an unusual mode of DNA interaction, with the binding interface formed by two HU dimers. The alpha helical region (AHR) of HU/IHF proteins consist an interface, which is laced with positive charged residues which we termed as DNA draping interface. DNA draping interface from two HU dimers interact with each other to form a DNA binding cradle (Fig 6). Other than conserved residue K3, we observed K18, which forms a part of HUpb interface and in E. coli interacts with GalR, also interacts with the DNA.

**Fig. 6:**
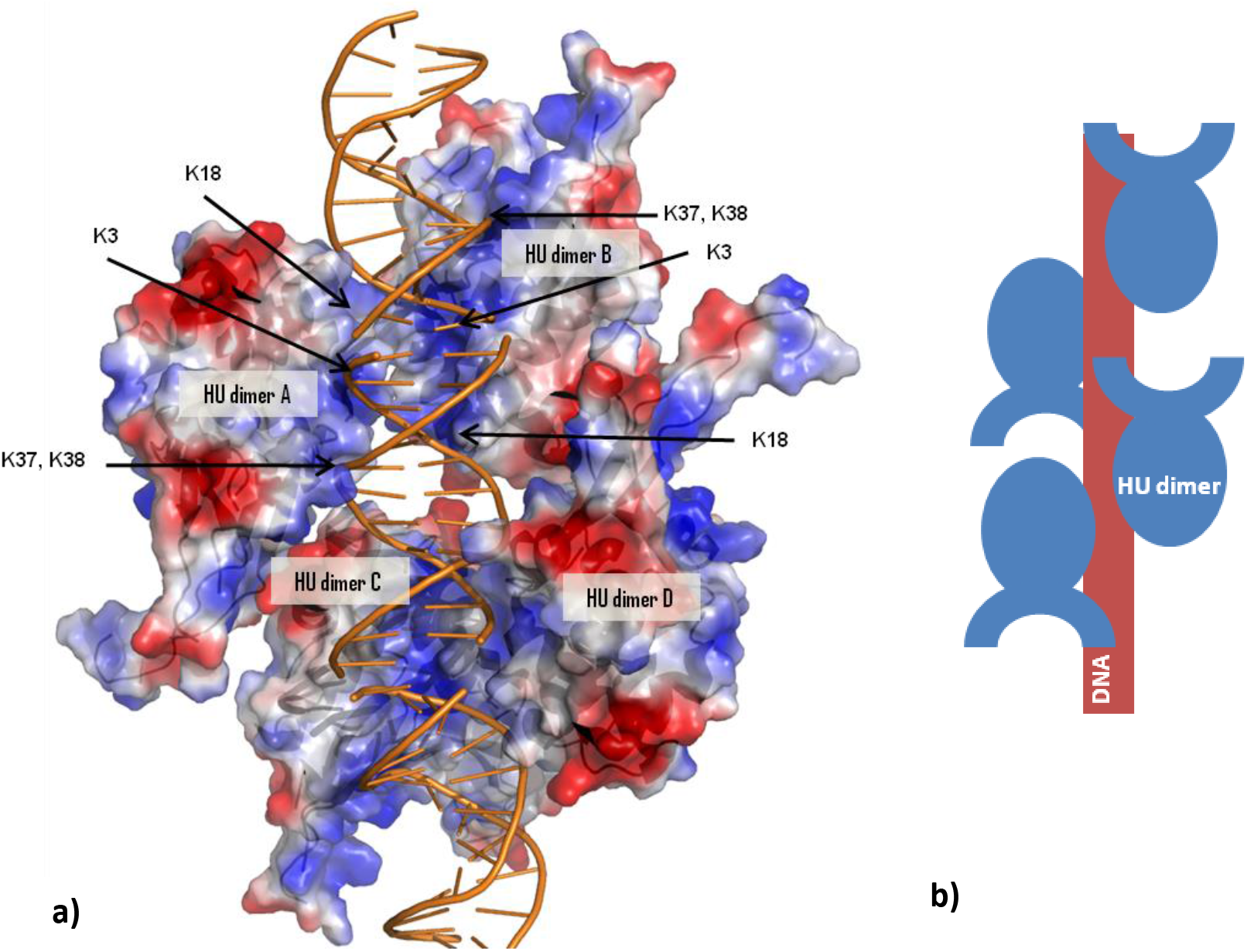
a) Supramolecular association of *E. coli* HU dimer (PDB ID: 4YF0) exhibiting an unusual mode of DNA interaction, with the binding interface formed by two HU dimers. The alpha helical region (AHR) of HU/IHF proteins consist an interface, which is laced with positive charged residues which we termed as DNA draping interface. The above arrangement makes use of the DNA draping interface from two HU dimers to interact with DNA. Other than conserved residue K3, we observed K18, which forms a part of HU_pb_ interface and in *E. coli* interacts with GalR, also interacts with the DNA, serving a dual purpose. Other residue which forms the DNA binding interface are K37 and K38. b) Schematic of HU-DNA interaction

Our observation is in full agreement with the functional data illustrating the role of residue position 17, 18 and 19 as hot-spot residues in protein-protein interface. Taken together, these data reveal that the evolution of residues at HUpb interface shows clade specific and phyla specific differences and in Proteobacteria, it can create various different permutations of interfaces in a single organism, which can differentially engage with different protein. Also, we found that the determinant of formation of pocket at the HUpb interface depends on the polarity of position 16.

### Reconstruction of ancestral HUpb interface residues

To gain insights into the evolutionary history of amino acid substitutions in HU/IHF family, we performed ancestral sequence reconstruction using ML method (Fig. 2). The ancestral sequences were extracted at the node prior to the HU/IHF clade division thus forming the root node (node 1). We also determined other ancestral sequences at the base of HU clade (node 2), IHFα clade (node 3), IHFβ clade (node4). We also investigated the ancestral sequences of the node prior to the divergence of Proteobacterial, Actinobacterial, Firmicutes HU (Node 5-7 respectively) and nodes at the base of sub-clades consisting African swine flu virus (node 8) and Spiroplasma melliferum (node 9). For each node, the best predicted sequence was analyzed. From the reconstruction, we were able to find the ancestral sequence prior to HU and IHF clade duplication event. Residue position 11, 12, 18 and 19 (w.r.t to MtbHU) are doesn’t determine the formation or absence of pocket.

Position 11 is majorly occupied by Ala, with all the sequences at the ancestral nodes (node 1-9) coding for the same, although we observe some characteristic differences in Actinobacteria and IHFα clade (Table 2). In MtbHU, this position is occupied by Thr while, in IHFα clade proteins, this position is aromatic in nature with Phe and Tyr as major residues. Position 12 is variable with differences within and among the HU-IHF clades. In Proteobacterial HUα sub-clade, negatively charged residues are favored while in Proteobacterial HUβ sub-clade, Ala is predominant (Table 2). Consistently, the same observation can be made for the HU homologs at the base of Proteobacterial HU node (node 5) occupied by Glu. We also observed that the root node (Node 1) along with HU and IHFα root nodes (Node 2 and 3 respectively) occupy Glu at this position. In Actinobacteria and Firmicutes, this position is favored by Glu and Gln (Q12 in MtbHU). While in Thermotogae Maritima, Lys is present at this position. In IHFα clade, position 12 is occupied mainly by Lys (in the Brucella genus) and Asp (in most of the γ-Proteobacteria). The root nodes at the base of sub-clade divisions in HU clade occupy Glu, thus making it a crucial conserved ancestral residue at this positon. As observed earlier, this position is negatively charged in both Firmicutes and γ-Proteobacterial IHFα (Table 1, Fig 5 c,i-k). Differences are observed in IHFβ clade, in which this position is variable with polar and charged residues making the major share (Table 2) and ancestral node for IHFβ at this position codes for Ser.

**Table 2:**
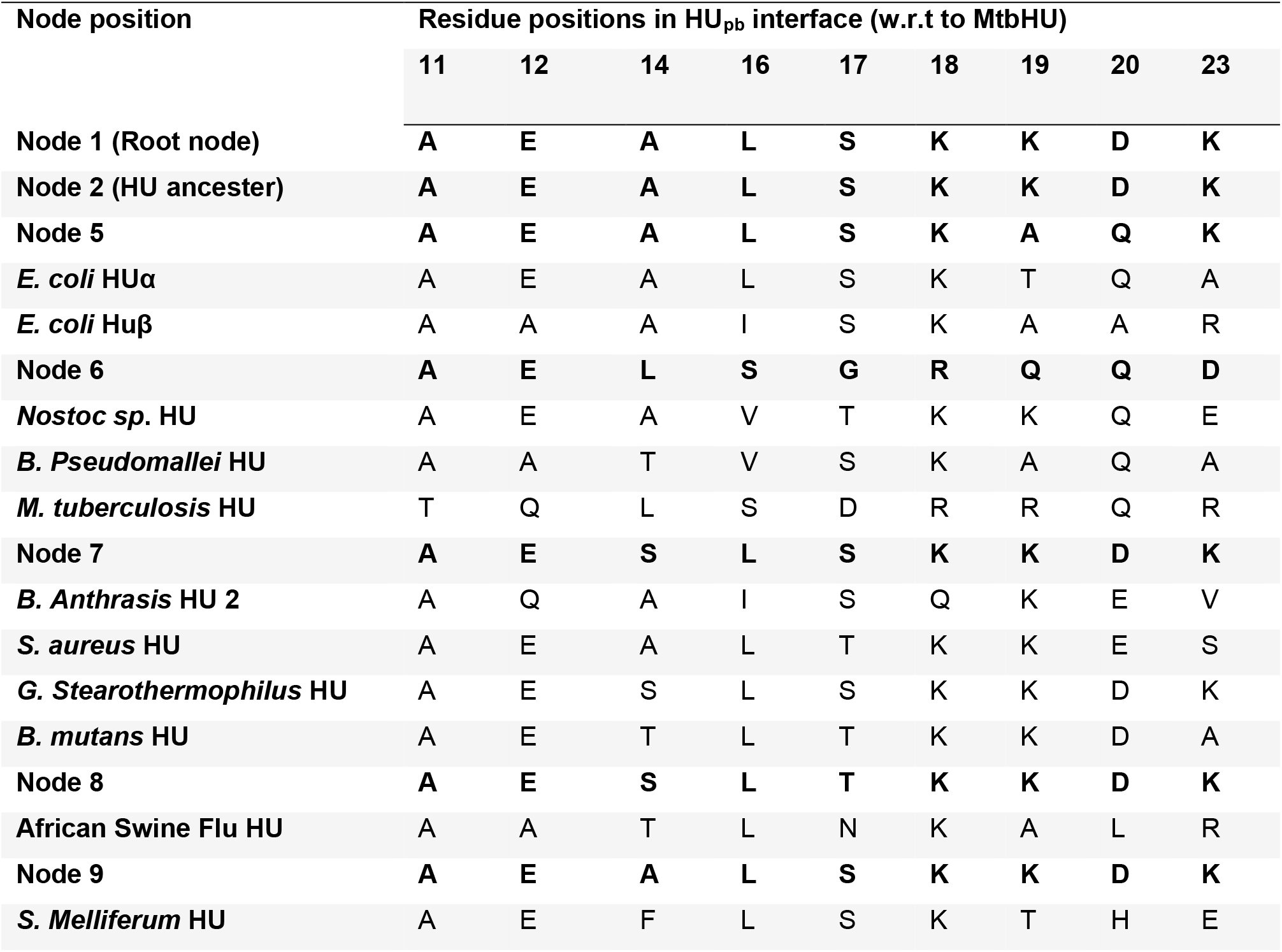
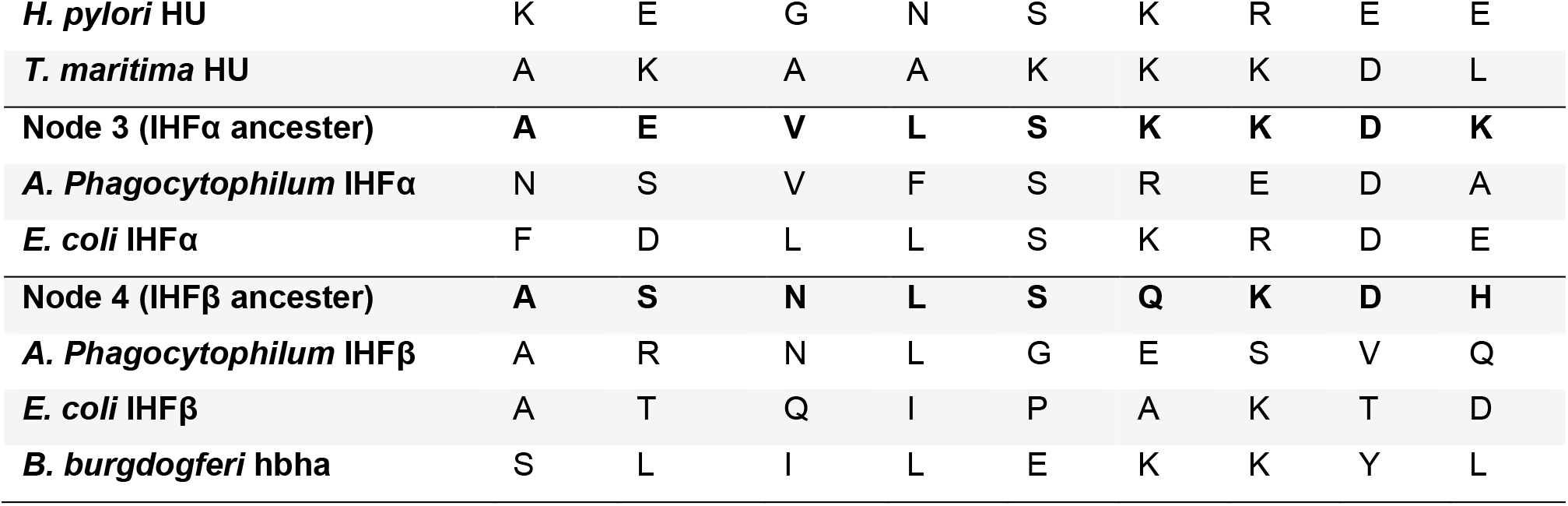
Residues at ancestral nodes and tips (representing HU and IHF proteins) at the HU_pb_ interface.

| Node position | Residue positions in HU<sub>pb</sub> interface (w.r.t to MtbHU) |  |  |  |  |  |  |  |  |
| --- | --- | --- | --- | --- | --- | --- | --- | --- | --- |
|  | 11 | 12 | 14 | 16 | 17 | 18 | 19 | 20 | 23 |
| Node 1 (Root node) | A | E | A | L | S | K | K | D | K |
| Node 2 (HU ancestor) | A | E | A | L | S | K | K | D | K |
| Node 5 | A | E | A | L | S | K | A | Q | K |
| *E. coli* HUα | A | E | A | L | S | K | T | Q | A |
| *E. coli* HUβ | A | A | A | I | S | K | A | A | R |
| Node 6 | A | E | L | S | G | R | Q | Q | D |
| *Nostoc sp.* HU | A | E | A | V | T | K | K | Q | E |
| *B. Pseudomallei* HU | A | A | T | V | S | K | A | Q | A |
| *M. tuberculosis* HU | T | Q | L | S | D | R | R | Q | R |
| Node 7 | A | E | S | L | S | K | K | D | K |
| *B. Anthraxis* HU 2 | A | Q | A | I | S | Q | K | E | V |
| *S. aureus* HU | A | E | A | L | T | K | K | E | S |
| *G. Stearothermophilus* HU | A | E | S | L | S | K | K | D | K |
| *B. mutans* HU | A | E | T | L | T | K | K | D | A |
| Node 8 | A | E | S | L | T | K | K | D | K |
| African Swine Flu HU | A | A | T | L | N | K | A | L | R |
| Node 9 | A | E | A | L | S | K | K | D | K |
| *S. Melliferum* HU | A | E | F | L | S | K | T | H | E |
| *H. pylori* HU | K | E | G | N | S | K | R | E | E |
| *T. maritima* HU | A | K | A | A | K | K | K | D | L |
| **Node 3 (IHFα ancestor)** | **A** | **E** | **V** | **L** | **S** | **K** | **K** | **D** | **K** |
| *A. Phagocytophilum* IHFα | N | S | V | F | S | R | E | D | A |
| *E. coli* IHFα | F | D | L | L | S | K | R | D | E |
| **Node 4 (IHFβ ancestor)** | **A** | **S** | **N** | **L** | **S** | **Q** | **K** | **D** | **H** |
| *A. Phagocytophilum* IHFβ | A | R | N | L | G | E | S | V | Q |
| *E. coli* IHFβ | A | T | Q | I | P | A | K | T | D |
| *B. burgdogferi* hbha | S | L | I | L | E | K | K | Y | L |

In MtbHU, we observed that L14 is one of the residues which form the pocket (Fig. 2b). This position is variable among HU and IHF clade proteins. In Proteobacterial HU, Ala is predominant, while in Actinobacteria it is occupied by Leu. At this position, polar residues Ser and Thr along with Ala is predominant in Firmicutes, while in Spiroplasma melliferum, it is occupied by Phe. In HU root node (Node 2) Ala is present which further got mutated to Leu at the base of the Actinobacterial HU root (Node 6). We also observed that it is mutated to polar residue Ser at the base of Firmicutes HU root (Node 7). At this position, hydrophobic residues (Leu, Ile and Val) are occupied by IHFα, proteins while it is occupied by polar residues in IHFβ clade proteins, which is reflected in their respective root nodes (Node 3and 4).

As previously discussed, residue position 16 plays a crucial role in the formation of pocket at the HUpb interface. Our phylogenetic and ancestral reconstruction data suggests that majorly this position is occupied by hydrophobic residues. In Mycobacterium genus it is occupied by Ser, while in Nocardia genus it is Thr (Table 2). We observed that the Actinobacterial root node (Node 6) is occupied by Ser; showing that a mutation at this site in the ancestral sequence became the determinant of pocket formation (Table 2). All the rest root nodes at the bases of dividing the clades and sub-clades consist Leu (Table 2).

Earlier studies have shown the role of residue position 17, in its playing key role in engaging the protein partner (Kar and Adhya 2001, Ghosh et al. 2015). From the phylogenetic tree we observed that in HU clade, it is majorly occupied by polar residues Ser and Thr whereas in Actinobacteria, it is occupied by either Gly or Asp (as in MtbHU D17, which interacts with Topoisomerase I with this residue). In IHFα clade proteins, this position is majorly occupied by Ser and Thr, similar to HU clade. In IHFβ clade, it is occupied majorly by hydrophobic residues. Phe is predominant in IHFβ proteins from Brucella and Bartonella genus at this position while most of the γ-Proteobacteria has Pro at this position. Our data found the root nodes at the bases of dividing the clades and sub-clades occupy mostly Se and Thr at this position (Table 2).

As discussed earlier, position 18 is mostly conserved with positively charge residue, might be playing dual role in interacting with proteins as well DNA. It is occupied majorly by Lys in HU and IHFα clade, with exception of Mycobacterium and Brucella genus, in which Arg is occupied. In our previous study, we already observed that Arg is majorly preferred in both BDR and AHR of Actinobacteria. In IHFβ clade, it is occupied majorly by polar residue Gln or Ala (in most of the γ-Proteobacteria). Although, in the next residue position (19), IHFβ clade is occupied by Lys, while IHFα clade is occupied by Arg (γ-Proteobacteria) and polar residues. At this position, HU clade exhibits sub-clade specific differences. Ala or polar residues occupy the Proteobacterial HU at this position, while positively charged residue Arg and Lys occupy this position in Actinobacterial and Firmicutes respectively (Table 2). Notebly, in IHFα clade, it is occupied by positively charged residues, but in IHFβ clade, it is majorly occupied by negatively charged or polar residues, with root node at the base of IHFβ clade, occupied by Gln(Table 2). We observed that position 19 is mostly positively charged and contributes to the DNA binding interface too, with most of its root node occupying Lys at this position (Table 2).

Another pocket lining residue position 20 (as per MtbHU), is observed to be majorly occupied by Asp or Gln (Table 2). Proteobacterial and Actinobacterial HU ancestral sequences were mutated at this node to Gln from Asp, which is the root node for HU (Node 2). It remained same for Firmicutes HU, and IHF clades, thus imparting another negatively charge residue in the HUpb interface. From our ancestral sequence reconstruction, we observed various mutational events at clade or sub-clade nodes which have both structural and functional consequences.

### Screening for inhibitors of MtbHUpb: A case study

As discussed in the previous section, other than the DNA binding interface (BDR), which we have previously inhibited using stillbene and suramin derivatives (Bhowmick et al. 2014), MtbHUpb interface is responsible for protein-protein interaction with DNA Topoisomerase I (Ghosh et al. 2015). Thus, it is an important interface which can be targeted with small molecules to disrupt MtbHU’s interaction with Topoisomerase I. Protein-protein interfaces are usually a challenging target due to its flatter surface. In case of MtbHUpb interface, we observed that due to the presence of a partially hydrophobic pocket, small molecule scaffolds could fit into it, while the ligand can be further grown to shield D17, which is the crucial residue for Topoisomerase I interaction (Ghosh et al. 2015). The following virtual screening, chemoinformatics analysis and iterative docking is performed on MtbHU as a case study, which can be further applied to other HU and IHF like proteins.

For identification of different inhibitory leads and their corresponding chemical scaffolds, we performed a two step virtual screening, using first the molecules listed in DrugBank database, aiming to repurpose existing inhibitors or their scaffolds. In discovering drug-like molecules for a myriad of diseases, repurposing of existing drugs can play a crucial role in speeding up the drug discovery pipeline (Opera and Mestres 2012). For known drugs, most of the toxicity, ADME and adverse effects are already known and tested, thus introducing the same or a slightly modified variant of that molecule is comparatively easier than a completely new molecule.

Docking of DrugBank molecules to MtbHUpb resulted ∼700 hits with docking score < −5 kcal/mol using High Throughput Virtual Screening (HTVS) of GLIDE, version 5.5 (Schrodinger, LLC, New York, 2009). Docking analysis with more precise XP GLIDE revealed 160 lead molecules with XP score less than −6.0 kcal/mol (Fig. 7). Our docking results showed compounds like Maltotetraose, Valrubicin, Iodixanol, Enalkiren, indinavir, Carfilzomib, oxytetracycline, quinalizarin, Raltitrexed, Epigallocatechin and their analogues exhibit high scoring binding with MtbHUpb pocket. Analyzing the interactions of MtbHUpb interface to ligands revealed some conserved as well as scaffold specific modes of interaction.

**Fig 7:**
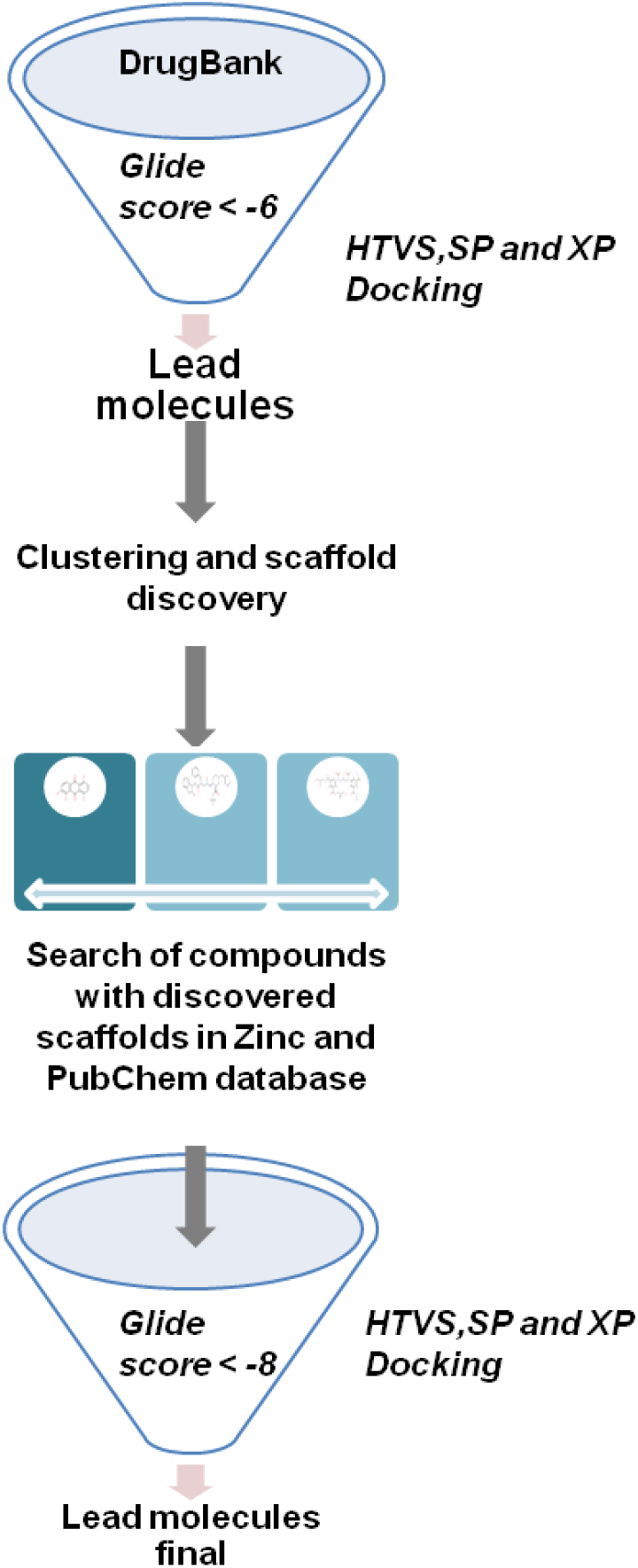
Two step virtual screening pipeline used in this study. Firstly, ligands of DrugBank were used for virtual screening and precision docking, which were further clustered to obtain key scaffolds. The chemical scaffold were then used as query for Zinc and PubChem database, for similar compounds which were again docked to obtain high scoring ligands.

Maltotetraose has the highest binding score (Glide XP score −10.38 kcal/mol) among DrugBank dataset, which interacts with L14 and Q20 of both chains via H-bond. It has a high number of rotatable bonds (10) and involves in H-bonding via hydroxyl group, while filling the MtbHUpb pocket completely (Fig. 8a). Although its binding score is high, it fails to interact or shield Asp17 residue, which is involved in Topoisomerase I interaction. Valrubicin is the next highest scoring (Glide XP score −9.31 kcal/mol) lead molecule, which interacts with Q20, D17, L14 from one chain and S16 from both the chains of MtbHU (Fig. 8b). It belongs to Anthracycline class of molecules which consists of a tetracenequinone ring linked with a sugar through a gycosidic linkage. The tetracenequinone ring is inserted into the bottom pocket, while the fluoride group attached to the sugar moiety interacts with D17. It belongs to the same class of molecules as Doxobubicin, which acts as a DNA binding drug used in cancer chemotherapy (Patel and Kaufmann 2012). In a similar way, Quinalizarin belonging to the same class (anthraquinones) fits into the MtbHUpb pocket and interacts to L14 of both the chains via polar hydroxyl groups (Fig. 8c).

**Fig 8:**
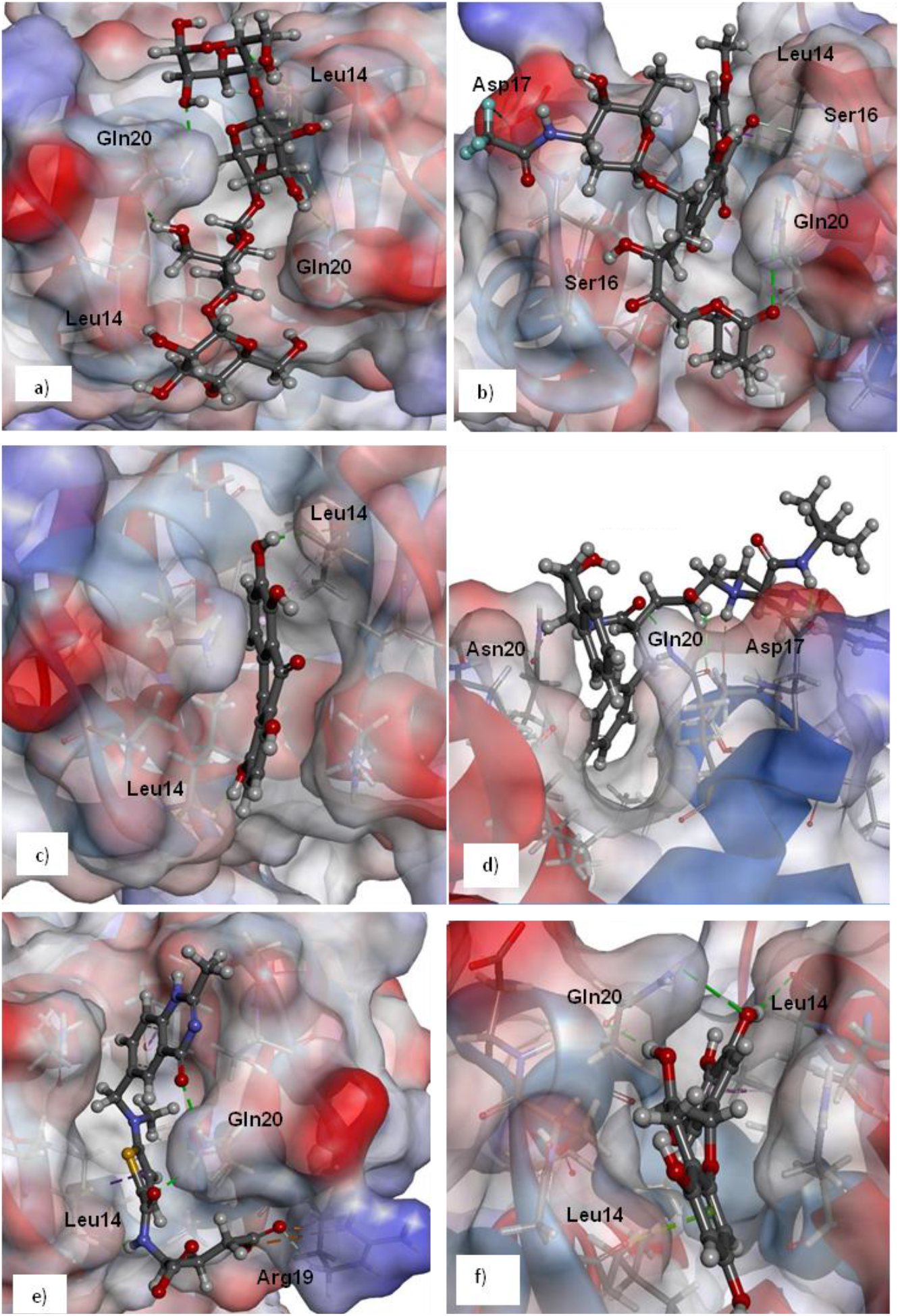
**First round of virtual screening and docking from DrugBank database** obtained ligands whose scaffolds were used for further rounds of docking. a) Maltotetraose, binding to the HU_pb_ pocket without obstructing Asp17. b) Valrubicin, which sits in the pocket and also shields Asp17. c) Quinalizarin, a small anthraquinone like compound which sits in the pocket interacting with L14. d) Indinavir sits in the pocket and also shields Asp17, while e) Raltitrexed shields Arg19. F) Epigallocatechin-3-gallate (EGCG), is seen to mainly inteact with pocket lining residues Leu14 and Gln 20

Our docking studies found Iodixanol although showing high docking score (Glide XP score −8.24 kcal/mol), interacts with only surface residues (Q20 and D17), without covering the pocket. Antiviral drug Indinavir along with other similar compounds, e.g. Saquinavir also shows promising docking scores and binding modes. Both the molecules and their analogues are HIV Protease inhibitors which are successfully used for AIDS therapy (Flexner 1998). It covers both with pocket and shields one of the D17 residues. In the pocket it interacts with Q20 from both the chains (Fig. 8d). We also observed that Raltitrexed, an inhibitor of thymidylate synthase binds with high docking score (−6.85 kcal/mol) and interacts with Q20, L14 and R19 which are the key residue for protein–protein interaction) (Fig. 8e). Quinalizarin belongs to the anthraquinones class of hydrophobic compounds which fits into the pocket and interacts to L14 of both the chains via polar hydroxyl groups (Fig. 8c). Epigallocatechinis are a flavonoid class of compounds, which recently has gained interest as antibiotic resistance breakers. Epigallocatechin-3-gallate (EGCG), is the major polyphenolic constituent present in green tea which inhibits grown and promote cell cycle apoptosis in Androgen-Sensitive and Androgen-Insensitive Human Prostate Carcinoma Cells (Gupta et al. 2000). It interacts with Q20 of one chain and both L14 residues with high docking score (−6.5 kcal/mol) (Fig. 8f). We clustered the molecules using hierarchical clustering using Canvas (Schrodinger, LLC, New York, 2009) to shortlist different scaffolds of the lead molecules to be used for further screening. The majority of the members of the cluster belong to anthraquinones followed by tetracenequinone analogues and flavenoid compounds. The core scaffold of the above listed compounds was given as query to search similar compounds (Tanimoto coefficient > 0.85) from PubChem and Zinc databases. We obtained > 5,00,000 compounds which were further docked using HTVS (High throughput), SP (Standard precision) and XP (Extreme precision) docking protocol in GLIDE, version 5.5 (Schrodinger, LLC, New York, 2009).

High scoring compounds binding to MtbHUpb interface were chosen which can fill up the MtbHUpb pocket, whilst shielding Asp17 (from at least one protomer chain). Our finding showed that tetracenequinone analogues which are similar to Valrubicin or Doxorubicin exhibited high docking scores (Table 4). The tetracenequinone ring sits into the hydrophobic pocket in a similar fashion in almost all the ligands in this category, while the flexible groups attached to the sugar moiety shields Asp17. We found tetracenequinone analog compound #1 (CID: 10349706) bind to MtbHUpb interface with docking XP score of −14.3 Kcal/mol. The tetracenequinone ring is inserted into the MtbHUpb pocket, where it is stabilized by Pi-alkyl interactions mediated by the aromatic rings of the molecule and alkyl groups of Leu14 and Ala23 (Fig. 9a). It is further stabilized by H-bonding interactions with pocket lining residues Leu 14 and Gln20 (Fig. 9a). The tetracenequinone ring is attached to a sugar moiety which is further extended as a poly-alcoholic chain. The hydroxyl groups of both the sugar moiety and alcoholic chain engage in H-bonding interaction with Asp17 (5.9a). Tetracenequinone analog compound #2 which is a isomer of #1 (CID: 44271259) bind to MtbHUpb interface in a slightly different manner, in which similar interactions are made by the tetracenequinone scaffold, while the extension makes extensive van der Waals as well as H-bonding interactions with Gln 20 and Asp17 (Fig. 9b). It binds to MtbHUpb interface with a docking XP score of −10.66 kcal/mol. Similar high scoring compounds with tetracenequinone scaffold (Compound #3-6) binding to MtbHUpb interface is listed in Table 4 and Fig. 9 c-f.

**Fig 9:**
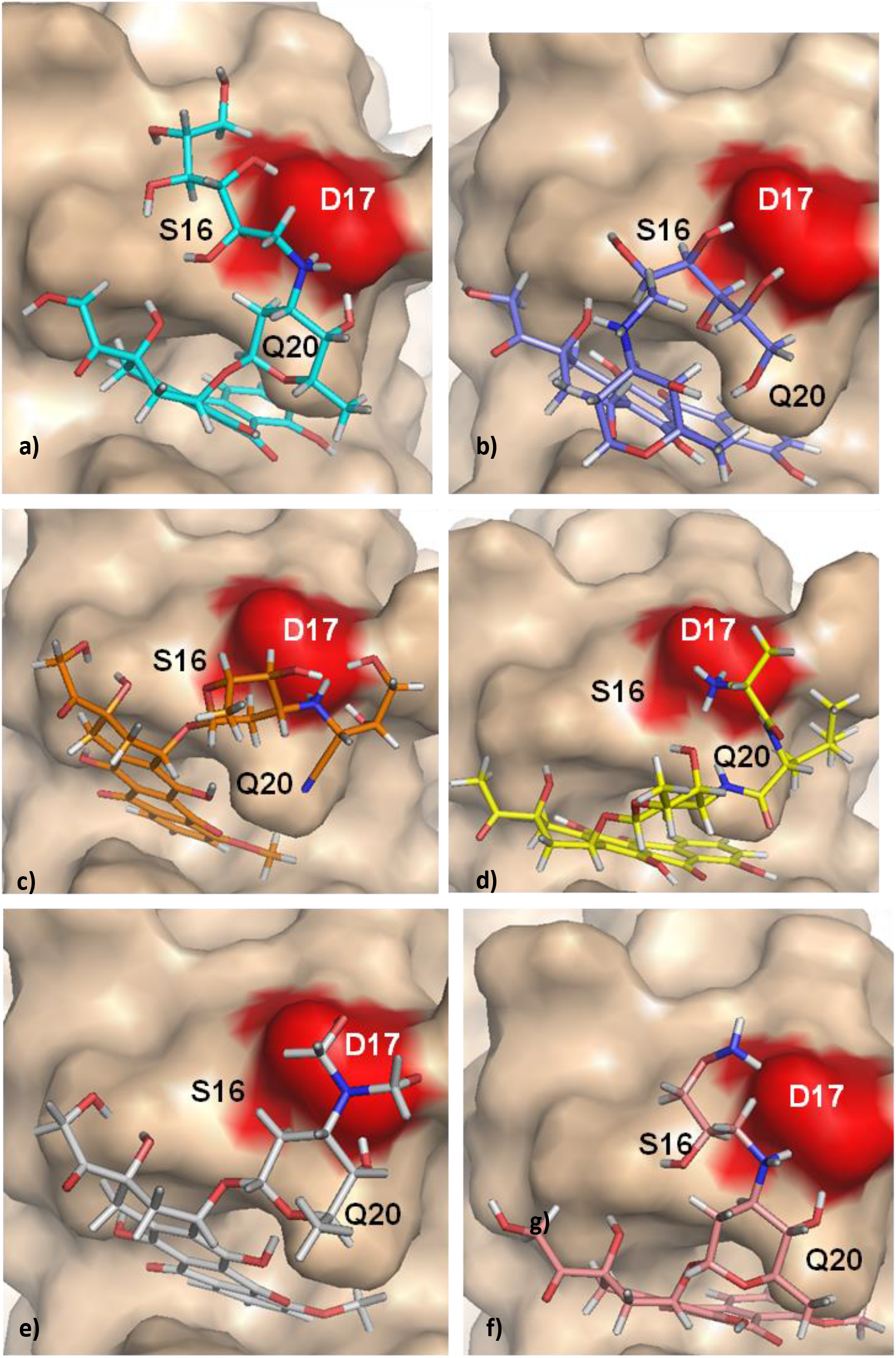
**a)-f)** Binding poses of ligand #1- 6 interacting with MtbHU_pb_ interface. All of the tetracenequinone analogs shows high scoring binding with MtbHU_pb_ interface, while interacting and shielding D17, which interacts with Topoisomerase I.

Other than tetracenequinone analogues, we also observed various polyphenolic compounds, which are analogues of indinavir and epigallocatechin (Table 4). Our scaffold based screening obtained a large variety of polyphenolic compounds which binds with high score to MtbHUpb interface. Compound #7 (CID: 72582243, XP docking score −9.3 Kcal/mol) partially sits into the MtbHUpb pocket, whilst a Pyrogallol moiety (aromatic ring with three hydroxyl substituent) interacts with Asp17 and Ser16 with H-bonding, shielding Asp17 (Fig 10a). Similar interactions are observed in compound #8 (CID: 59867087, XP docking score −9.3 Kcal/mol), #9 (CID: 442682, XP docking score −8.5 Kcal/mol), and #11 (CID: 5089687, XP docking score −9.4 Kcal/mol) in which the Pyrogallol moiety is used for shielding the Asp17 neighborhood(Fig. 10 b,c,e).

**Fig 10:**
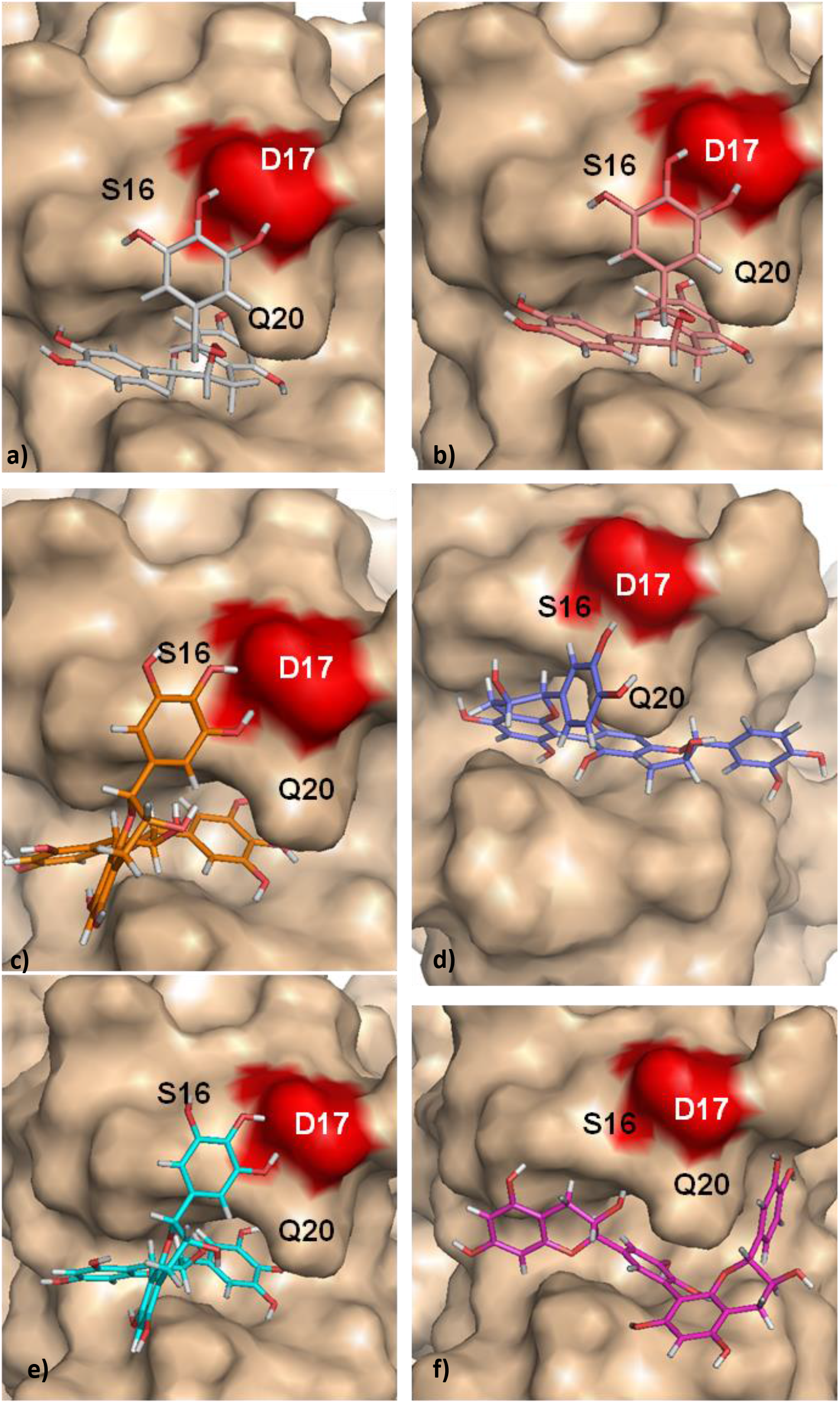
**a)-f)** Binding poses of ligand #7- 12 interacting with MtbHU_pb_ interface. All of the polyphenol analogs shows high scoring binding with MtbHU_pb_ interface, while a)- e) interacting and shielding D17, which interacts with Topoisomerase I.

Compound #10 (CID: 9808324, XP docking score −8.2 Kcal/mol) forms a more longer scaffold which fits into the MtbHUpb pocket, interacting with residues Gln20 and Ala23 from both chains. A Pyrocatechol moiety in this molecule shields the Asp 17 and Gln20, making H-boding interaction with them and further stabilized by Pi-alkyl interactions with Leu16 (Fig 10d). A similar compound #12 (CID: 5089687, XP docking score −8.8 Kcal/mol), composed of Pyrocatechol moiety, was observed to extensively fill the MtbHUpb pocket, without making any interaction to Asp17(Fig 10f). Although having a higher score than compound #10, it doesn’t shield Asp17, thus can be taken as control ligand to understand the role of Asp17 shielding in disruption of MtbHU-Topoisomerase I interactions.

## Conclusion

This work gives a model example of an evolutionary study of a protein interface of nucleoid associated protein, which is used to understand the interface and computationally design inhibitors targeting it. This strategy could be in general useful for designing inhibitors all types of protein-protein interfaces, where evolutionary studies can guide to direct which interfaces are difficult (flatter interface) or which are easier (interface with pocket, which can act as an anchor for the core of inhibitory compound) to target.

## References

Abascal, F., Zardoya, R. and Posada, D., 2005. ProtTest: selection of best-fit models of protein evolution. Bioinformatics, 21(9), pp.2104–2105.

Bensaid, A., Almeida, A., Drlica, K. and Rouviere-Yaniv, J., 1996. Cross-talk Between Topoisomerase I and HU inEscherichia coli. Journal of molecular biology, 256(2), pp.292–300.

Bhowmick, T., Ghosh, S., Dixit, K., Ganesan, V., Ramagopal, U.A., Dey, D., Sarma, S.P., Ramakumar, S. and Nagaraja, V., 2014. Targeting Mycobacterium tuberculosis nucleoid-associated protein HU with structure-based inhibitors. Nature communications.

Dey, D., Nagaraja, V., & Ramakumar, S. (2017). Structural and evolutionary analyses reveal determinants of DNA binding specificities of nucleoid-associated proteins HU and IHF. Molecular phylogenetics and evolution, 107, 356–366.

Dey, Debayan. “Crystal Structures of Native and AdoMet Bound rRNA Methyltransferase from Sinorhizobium meliloti: Structural Insights into rRNA Recognition. Evolutionary, Structural and Functional Studies on Nucleoid-Associated Proteins HU and IHF.”. PhD diss., 2018.

Dey, D., Kavanaugh, L. G., & Conn, G. L. (2020). Antibiotic substrate selectivity of Pseudomonas aeruginosa MexY and MexB efflux systems is determined by a Goldilocks affinity. Antimicrobial Agents and Chemotherapy.

Dri, A.M., Rouviere-Yaniv, J. and Moreau, P.L., 1991. Inhibition of cell division in hupA hupB mutant bacteria lacking HU protein. Journal of bacteriology, 173(9), pp.2852–2863.

Flexner, C. 1998. HIV-protease inhibitors. N Engl J Med, 338(18), 1281–1292.

Ghosh S, Mallick B, Nagaraja V. 2015. Direct regulation of topoisomerase activity by a nucleoid-associated protein. Nucleic acids research, 42(17): 11156–11165.

Giladi, H., Koby, S., Prag, G., Engelhorn, M., Geiselmann, J., & Oppenheim, A. B. 1998. Participation of IHF and a distant UP element in the stimulation of the phage λ PL promoter. Molecular microbiology, 30(2), 443–451.

Goh, C.S., Bogan, A.A., Joachimiak, M., Walther, D. and Cohen, F.E., 2000. Co-evolution of proteins with their interaction partners. Journal of molecular biology, 299(2), pp.283–293.

Gupta, S., Ahmad, N., Nieminen, A.L. and Mukhtar, H., 2000. Growth inhibition, cell-cycle dysregulation, and induction of apoptosis by green tea constituent (-)-epigallocatechin-3-gallate in androgen-sensitive and androgen-insensitive human prostate carcinoma cells. Toxicology and applied pharmacology, 164(1), pp.82–90.

Hwang, D. S., & Kornberg, A. (1992). Opening of the replication origin of Escherichia coli DnaA protein with protein HU or IHF. Journal of Biological Chemistry, 267(32), 23083–23086.

Jayaraman, L., Moorthy, N.C., Murthy, K.G., Manley, J.L., Bustin, M. and Prives, C., 1998. High mobility group protein-1 (HMG-1) is a unique activator of p53. Genes & development, 12(4), pp.462–472.

Jeffery CJ. Moonlighting proteins: old proteins learning new tricks. TRENDS in Genetics. 2003 Aug 31;19(8):415–7.

Kamashev, D. and Rouviere-Yaniv, J., 2000. The histone-like protein HU binds specifically to DNA recombination and repair intermediates. The EMBO journal, 19(23), pp.6527–6535.

Kar S, Adhya S. 2001. Recruitment of HU by piggyback: a special role of GalR in repressosome assembly. Genes & development, 15(17): 2273–2281.

Katsube, T., Matsumoto, S., Takatsuka, M., Okuyama, M., Ozeki, Y., Naito, M., Nishiuchi, Y., Fujiwara, N., Yoshimura, M., Tsuboi, T. and Torii, M., 2007. Control of cell wall assembly by a histone-like protein in mycobacteria.Journal of bacteriology, 189(22), pp.8241–8249.

Khare, H., Dey, D., Madhu, C., Senapati, D., Raghothama, S., Govindaraju, T., & Ramakumar, S. (2017). Conformational heterogeneity in tails of DNA-binding proteins is augmented by proline containing repeats. Molecular BioSystems, 13(12), 2531–2544.

Kuiper, E. G., Dey, D., LaMore, P. A., Owings, J. P., Prezioso, S. M., Goldberg, J. B., & Conn, G. L. (2019). Substrate recognition by the Pseudomonas aeruginosa EF-Tu–modifying methyltransferase EftM. Journal of Biological Chemistry, 294(52), 20109–20121.

Merika, M. and Orkin, S.H., 1995. Functional synergy and physical interactions of the erythroid transcription factor GATA-1 with the Krüppel family proteins Sp1 and EKLF. Molecular and cellular biology, 15(5), pp.2437–2447.

Le, S.Q. and Gascuel, O., 2008. An improved general amino acid replacement matrix. Molecular biology and evolution, 25(7), pp.1307–1320.

Oprea, T.I. and Mestres, J., 2012. Drug repurposing: far beyond new targets for old drugs. The AAPS journal, 14(4), pp.759–763.

Onrust, S. V., & Lamb, H. M. 1999. Valrubicin. Drugs & aging, 15(1), 69–75

Phan, J., Koli, S., Minor, W., Dunlap, R. B., Berger, S. H., & Lebioda, L. 2001. Human thymidylate synthase is in the closed conformation when complexed with dUMP and raltitrexed, an antifolate drug. Biochemistry,40(7), 1897–1902

Patel, A.G. and Kaufmann, S.H., 2012. How does doxorubicin work?. Elife,1, p.e00387.

Ryan, V.T., Grimwade, J.E., Nievera, C.J. and Leonard, A.C., 2002. IHF and HU stimulate assembly of pre-replication complexes at Escherichia coli oriC by two different mechanisms. Molecular microbiology, 46(1), pp.113–124.

Shrilakshmi, S., Kundapura, S., Dey, D., Ramagopal, U. A., & Kulal, A. (2019). Phylogenetic, sequence and structural analysis of Insulin superfamily proteins reveals an indelible link between evolution and structure-function relationship. bioRxiv, 769497.

Thomas, J.O. and Travers, A.A., 2001. HMG1 and 2, and related ‘architectural’DNA-binding proteins. Trends in biochemical sciences, 26(3), pp.167–174.

Tina, K.G., Bhadra, R. and Srinivasan, N., 2007. PIC: protein interactions calculator. Nucleic acids research, 35(suppl 2), pp.W473–W476.

van Noort, J., Verbrugge, S., Goosen, N., Dekker, C. and Dame, R.T., 2004. Dual architectural roles of HU: formation of flexible hinges and rigid filaments. Proceedings of the National Academy of Sciences of the United States of America, 101(18), pp.6969–6974.

Zappavigna, V., Falciola, L., Helmer-Citterich, M., Mavilio, F. and Bianchi, M.E., 1996. HMG1 interacts with HOX proteins and enhances their DNA binding and transcriptional activation. The EMBO Journal, 15(18), p.4981.

